# Post-admixture selection favours Duffy negativity in the Lower Okavango Basin

**DOI:** 10.1101/2025.11.06.687002

**Authors:** Artem Kasianov, Anne-Maria Fehn, Maitseo Bolaane, Gaseitsiwe Masunga, Ezequiel Chimbioputo Fabiano, Jorge Rocha

## Abstract

The FY*B^ES^ allele in the human Duffy blood group is nearly fixed across much of sub-Saharan Africa. Individuals homozygous for this allele (Duffy-negative) were considered resistant to red blood cell invasion by the malaria parasite *Plasmodium vivax*, restricting its distribution across the continent. However, as recent studies have demonstrated that *P. vivax* can infect Duffy-negative individuals and is widespread among African populations where FY*B^ES^ predominates, long-standing assumptions about the evolutionary relationship between Duffy negativity and parasite resistance should be re-evaluated across diverse geographic, ecological and epidemiological settings. Previous research investigating the role of natural selection in increasing the frequency of the FY*B^ES^ allele has primarily centered on admixed populations with African and non-African ancestries from regions with long-documented *P. vivax* transmission. Here, we focus on the Khwe foragers from the lower Okavango River Basin, where the parasite has only recently been reported. Using locus-specific statistics, simulations of neutral scenarios, and local ancestry inference, we found strong evidence for post-admixture selection promoting FY*B^ES^ introgression from Bantu-speaking populations into the Khwe. If *P. vivax* resistance indeed drove the rise in FY*B^ES^ allele frequency, our findings suggest that the parasite has been present in the region at least 500 to 1,000 years ago.

## 1. Introduction

The human Duffy blood group (FY) antigens act as chemokine receptors and facilitate the invasion of red blood cells (RBC) by the malaria parasite *Plasmodium vivax* [1–3]. Most antigenic variation in FY is determined by three common alleles—FY*A, FY*B, and FY*B^ES^ —at the Atypical Chemokine Receptor 1 (ACKR1) gene on chromosome 1, also known as the Duffy Antigen Receptor for Chemokines (DARC) [4,5].

The FY*A (rs12075*G) and FY*B (rs12075*A) alleles synthesize functional transmembrane glycoproteins and are widespread in Eurasia and the Americas. FY*B^ES^ is an erythroid-silent (ES) FY*B variant that is nearly fixed in much of sub-Saharan Africa and carries an additional promoter mutation (rs2814778 T>C), which leads to the loss of FY antigen expression in red blood cells (RBC) [6–9].

Because homozygosity for the FY*B^ES^ allele (Duffy-negativity) was long thought to completely block *P. vivax* invasion of RBC, it has been hypothesized that selective pressure from *vivax* malaria caused the fixation of the allele, thus limiting the spread of the parasite across large regions of Africa due to the lack of susceptible hosts [1,10,11].

More recently, however, molecular diagnostic tools have revealed that *P. vivax* can infect Duffy-negative individuals and is widespread across African populations where Duffy-negativity is near fixation [12–14]. Still, as infections appear less severe in Duffy-negative than in Duffy-positive individuals [14–17], it remains unclear to what extent this finding challenges the hypothesis that *P. vivax* was an important selective force driving the spread of Duffy-negativity.

To address this question, it is important to examine the impact of natural selection on FY genotypes across widely distributed geographic, ecological, and epidemiological contexts. So far, signatures of ongoing positive selection increasing FY*B^ES^ allele frequency have only been described in admixed populations of African and non-African ancestry, in areas where *P. vivax* has been historically prevalent [18–22].

Here we investigate the role of natural selection in elevating the frequency of the FY*B^ES^ allele in the lower Okavango River Basin, an African region where the parasite was only recently found to coexist with high frequencies of Duffy-negativity [23,24].

By combining locus-specific and local ancestry approaches using genome-wide data, we found strong evidence that post-admixture selection has promoted the diffusion of the FY*B^ES^ allele from Bantu-speaking populations into the Khoe-Kwadi–speaking Khwe, a foraging group, that arose from admixture between resident southern African hunter-gatherers, migrant herders from East Africa and Bantu-speaking agriculturalists originating in West Africa [25,26].

If *P. vivax* resistance is indeed responsible for the rise in FY*B^ES^ allele frequency, our findings indicate that the parasite was present in the region at least 500 to 1,000 years ago.

## 2. Material and Methods

### 2.1 Sample information

This study includes 149 newly genotyped samples from two ethnolinguistic groups in Namibia and Botswana: the Khwe (114 individuals) and the Kwangali (35 individuals) (Fig. S1; Table S1).

The Khwe speak a Khoe-Kwadi language [27] and have traditionally subsisted on foraging in areas of the Okavango River Basin spanning across southeastern Angola, northeastern Namibia, and Botswana’s Okavango Delta [28]. Based on linguistic variation and livelihood patterns, the Khwe can be divided into two main subgroups, both of which are represented in our sample: the □Ani-Khwe (30 individuals), primarily fishermen currently residing near the Okavango Delta in northwestern Botswana; and the non-□Ani-Khwe (84 individuals), who speak closely related □Xom, □Xoo, and Buga varieties, and historically inhabited the drier regions to the east and north of the Okavango Delta [29] (Figs. S1 and S2). The Kwangali are a West Bantu-speaking group within the Kavango language cluster (K.33), who primarily practice agriculture along the Namibian and Angolan banks of the lower Okavango River [30].

All data were collected under the TwinLab collaborative network linking CIBIO/InBIO with the University of Namibia and the University of Botswana. Official research permits were granted by the Ministries of Health of Namibia (RPIV00722019) and Botswana (HPRD: 6/14/1). At each sampling site, the study was explained to participants in their native language by a bilingual translator. Written informed consent was obtained from all volunteers prior to sample collection. From each participant, we collected a saliva sample along with ethnolinguistic information, including ethnic affiliation, place of birth, and language spoken up to the grandparental generation.

### 2.2 Genotyping and quality control

We genotyped all sampled individuals using the Affymetrix Axiom Genome-Wide Human Origins Array [31], and performed quality control and filtering with PLINK v1.9 [32].

Non-biallelic single nucleotide polymorphisms (SNPs), SNPs with call error rates exceeding 10%, and non-autosomal SNPs were excluded. After filtering, a total of 553,841 SNPs remained. No individuals had missing rates greater than 10%.

For comparative purposes, we merged our data with that of 130 individuals across 14 contextual African populations previously genotyped on the same array, also using PLINK v1.9 (Fig. 1; Table S1). The intersection between the two datasets yielded a total of 535,116 SNPs.

**Fig. 1.**
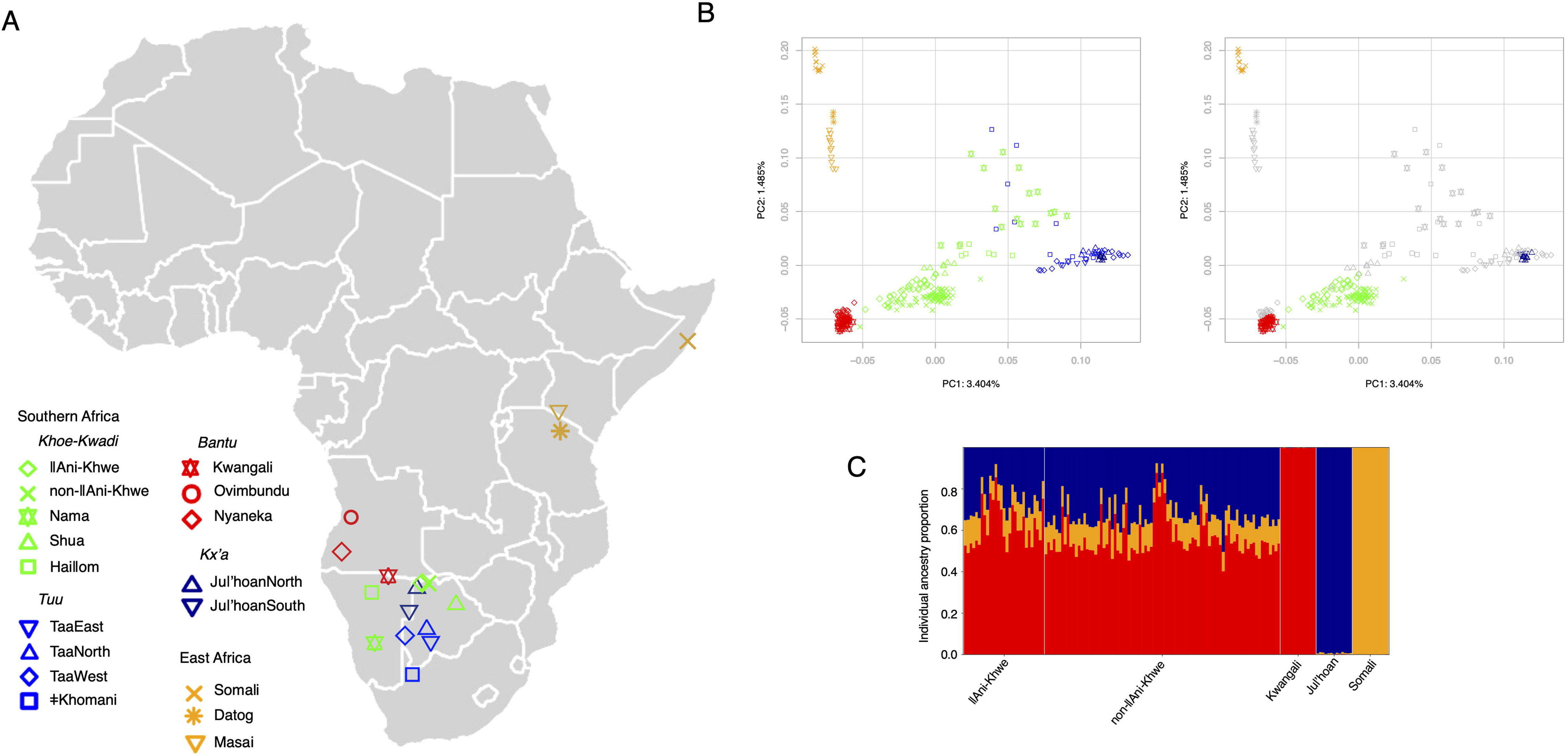
Population structure and admixture in the Khwe. (A) Map showing the approximate geographic locations of newly sampled and contextual populations. (B) PCA plots of the populations shown in the map. In the right panel, only Khwe individuals and parental populations are highlighted in colour. (C) Individual ancestry proportions inferred using RFMix v2. Vertical lines represent the estimated ancestry proportions for each individual.

To independently genotype SNPs rs2814778 and rs12075, which determine Duffy allelic variation, we used Sanger sequencing following polymerase chain reaction (PCR) amplification with previously described primers [7].

### 2.3 Phasing

Phasing was performed with Beagle 5.4 [33], employing the standard HapMap genetic map available on the Beagle website (https://faculty.washington.edu/ browning/ beagle/beagle.html).

To ensure results were not dependent on a specific phasing method, we also conducted analyses with SHAPEIT v2.r904 [34], using the HapMap b37 genetic map available on GitHub (https://github.com/odelaneau/shapeit4/tree/master/maps).

### 2.4 Population structure analysis

To examine population structure, we performed principal component analysis (PCA) using a dataset pruned for linkage disequilibrium (LD) with PLINK v1.9 [32]. The pruned dataset comprises 343,959 SNPs and was generated by removing SNPs with an r² value greater than 0.4 within 200-kb windows, shifted at 25 SNP intervals. Principal component scores were computed with PLINK and visualized with a *PlotPCAs.r* script.

### 2.5 Ancestry inference

Ancestry inference at the population, individual and local-levels was performed using RFMix v2 [35] with phased samples and a three-way admixture model, assuming an admixture time of 25 generations. The analysis included 114 Khwe individuals and employed two reference panels with samples of equal size, representing ancestries associated with the Bantu expansion, the indigenous southern African population layer, and the eastern African pastoral migration. The first panel consisted of 13 Kwangali (Bantu ancestry), 13 Ju□’hoan (southern African hunter-gatherers), and 13 Somali (eastern African pastoralists) individuals. In the second panel, the Somali samples were replaced with 15 Hadza individuals to assess the impact of using different Eastern African sources on local ancestry profiles; the sample sizes for the Kwangali and Ju□’hoan were increased accordingly.

We used a two SNP window size, three Expectation-Maximization iterations and the option “reanalyze-reference” to account for possible admixture in the reference groups. To test the robustness of our results, we used an alternative admixture time of 39 generations corresponding to the average between proposed dates of 60 generations ago for the initial admixture between East Africans and indigenous southern Africans, and 17 generations ago for the later admixture with Bantu-speaking groups [25].

Individual ancestry proportions were calculated by averaging the chromosome-specific ancestry estimates provided in RFMix *Q* output files and were visualized using a Python script (*VisualizeAdmixture.rfmix.py*). Population-level ancestry was then determined by averaging the individual ancestry proportions within each group.

Local ancestry assignments were performed using the RFMix *fb* output files, which provide marginal probabilities for each genomic window originating from a given parental population. Windows with marginal probabilities less than 1 were excluded to remove segments with ambiguous ancestry assignments. In addition, we filtered out SNPs located in centromeric regions and within 2 Mb of chromosome telomeres [36–38].

### 2.6 Assessing deviations from expected admixture allele frequencies

To evaluate whether the frequency of FY*B^ES^ in the Khwe population was higher than expected from admixture alone, we employed a binomial resampling approach, where the probability of success was defined as the product of the FY*B^ES^ allele frequency in the ancestral populations and their estimated admixture proportions [39]. The resampling procedure is implemented in the *resampling.CLI.plots.py* script.

Additionally, we used the F_adm_ statistic [40] to compare the deviations from expected admixture allele frequencies at the Duffy locus with those observed at other SNPs genotyped on the Genome-Wide Human Origins Array.

### 2.7 Simulations of genetic drift

We performed forward simulations under the Wright–Fisher model to generate expected distributions of FY*B^ES^ allele frequencies in the absence of selection, using the *Wright_Fisher_simulator.py* script. The simulated admixed population was modelled as initially formed through contributions from resident southern African hunter-gatherers and incoming East African herders, followed by a secondary admixture event involving Bantu-speaking populations. We investigated the outcomes of varying effective population sizes, parental allele frequencies, admixture dynamics, and number of generations since admixture on the resulting allele frequency distributions.

### 2.8 Estimation of selection coefficient

We applied a deterministic framework, using equation 10.25 from [41] to estimate selection coefficients (*s*) that could account for the observed FY*B^ES^ allele frequencies, given different initial allele frequencies and varying number of generations since admixture. Allele frequency trajectories were calculated across a range of *s* values from 0 to 0.4, in increments of 0.001, using dominance coefficients (*h*) set at 1 (recessive model) or 0.5 (additive model). The selection coefficient that yielded the FY*B^ES^ allele frequency closest to the observed value was taken as the best estimate. The procedure was implemented using the *Selection_coefficient_estimator.py* script.

## 3. Results

### 3.1 The admixture structure of the Khwe population

We assessed the population structure of the Khwe in comparison to other Khoe-Kwadi–speaking groups, as well as selected populations representing resident southern African, eastern African, and Bantu-related ancestries (Fig. 1). In PCA, the Khwe are positioned between Bantu-speaking groups and Kx’a- and Tuu-speaking southern African hunter-gatherers (PC1 in Fig. 1B), while also showing slightly greater proximity to eastern Africans than the Bantu populations included in this study (PC2 in Fig. 1B).

Previous studies have shown that the genetic composition of the Khwe and other Khoe-Kwadi–speaking populations can be modelled as the result of an initial admixture between East African and indigenous southern African–related ancestries, followed by a second admixture event with incoming Bantu-speaking groups [25]. Assuming a generation time of 30 years [42], inferred dates for these events range from approximately 2,100–1,800 years ago (∼70–60 generations ago) for the admixture between southern and easter African ancestries, and from 1,000–500 years ago (∼33–17 generations ago) for the admixture between the pre-Bantu hybrid population and the Bantu migrants [25,26,43,44].

Using RFMixv2 [35] and selecting the Kwangali, Ju□’hoan, and Somali as proxies for the three parental populations, we estimated that Khwe individuals carry, on average, 58.0% Bantu-related ancestry (standard deviation (SD) 9.8%), 30.5% (SD 8.6%) resident southern African–related ancestry, and 11.5% (SD 2.6%) eastern African–related ancestry (Fig. 1C; Table S2).

Our choice of parental populations was based on geographic, historical, and genetic criteria (Fig. 1A; Fig. S1). The West Bantu–speaking Kwangali, who inhabit the middle reaches of the Okavango River Basin, and the Kx’a-speaking Ju□’hoan, who live to the west of the Okavango Delta, both show strong genetic affinities with groups that have a documented history of contact and admixture with the Khwe [26,45–47]. In addition, the Afro-Asiatic–speaking Somali share approximately 62% of their genome with a ∼3,100-year-old individual from the Luxmanda pastoralist site in Tanzania, whose ancestry is closely related to the eastern African component found in modern Khoe-Kwadi–speaking populations [26,48].

As Khwe-speakers fall into two broad ethnolinguistic subgroups with slightly different genetic profiles (Fig. 1B; Fig. S2), here referred to as □Ani-Khwe and non-□Ani-Khwe, we additionally report results for these two sample subsets. On average, □Ani-Khwe individuals show higher Bantu-related ancestry (62.6% vs. 56.3%) and lower resident southern African–related ancestry (25.5% vs. 32.2%) compared to non-□Ani-Khwe individuals (Table S2). Despite these differences, the overall ancestry profiles are consistent with previous estimates—obtained with different methods and smaller sample sizes— showing that the Khwe carry a higher proportion of Bantu than resident southern African– related ancestry [25,26] (Table S2).

### 3.2 Elevated FY*B^ES^ frequency in the Khwe

We next investigated whether the frequency of FY*B^ES^ in the Khwe was higher than expected from admixture alone [39,49].

Because the SNPs rs12075 and rs2814778, which determine Duffy allelic variation, are not included in the Affymetrix Axiom Genome-Wide Human Origins array, we directly genotyped these two polymorphisms in the 114 Khwe and 26 Kwangali (Table S3). For the Somali and Ju□’hoan populations, we used previously published FY*B^ES^ allele frequencies [50,51] (Table S4).

Using a binomial resampling procedure with 10□replicates, we found that the observed FY*B^ES^ frequency of 94% in the Khwe is significantly higher (p < 10□□) than the 72% expected on the basis of their estimated ancestry proportions (Table S2) and the current FY*B^ES^ allele frequencies in the Kwangali (100%), Ju□’hoan (11%), and Somali (92%) source populations (Fig. S3; Table S4).

This approach is conservative, as the FY*B^ES^ allele was likely introduced into the Ju□’hoan and other resident southern African populations only recently, through admixture with Bantu-speaking groups [52]. A similar bias may result from using present-day FY*B^ES^ frequencies in eastern Africa to infer allele frequencies in the migrating pastoralists who initially reached southern Africa around 2,000 years ago [46]. If FY*B^ES^ entered eastern Africa during the Bantu expansions – and considering genetic evidence that admixture between Bantu-speaking migrants and resident eastern African populations occurred as recently as ∼1,000 years ago [44,48,53] – it is plausible that eastern African pastoralists migrated southward before acquiring the FY*B^ES^ allele through contact with Bantu speakers. Under this scenario, the pre-Bantu hybrid population in southern Africa would have entirely lacked the FY*B^ES^ allele, and its expected frequency in the Khwe following Bantu admixture would be close to 58% rather than 72%—further increasing the discrepancy with the observed frequency of 94%.

To further evaluate this deviation, we assessed the genome-wide distribution of the per-SNP F_adm_ statistic [40] to determine whether the divergence at the Duffy locus was unusually extreme compared to other loci. Notably, only 0.03% of all SNPs exhibited F_adm_ values equal to or greater than that of rs2814778, the SNP defining the FY*B^ES^ allele—even under the most conservative assumptions for parental allele frequencies: Kwangali 100%, Ju□’hoan 11%, and Somali 92% (Fig. S4). In a less conservative scenario, assuming that the FY*B^ES^ allele was absent in the pre-Bantu hybrid population, rs2814778 displayed the highest genome-wide F_adm_ value (Fig. S4).

Because the effects of genetic drift are expected to be distributed across the genome, while signatures of positive selection are typically localized around the selected locus, the outlier status of the Duffy locus suggests that the elevated frequency of FY*B^ES^ is unlikely to be explained by neutral evolutionary processes alone.

### 3.3 Effect of genetic drift

To further assess whether the elevated frequency of FY*B^ES^ in the Khwe could be explained by genetic drift, we conducted simulations under a Wright–Fisher model. We explored a general framework in which a pre-Bantu population is first formed through a single pulse of admixture between southern African foragers and eastern African herders, followed by subsequent genetic contributions from Bantu-speaking migrants (Fig. 2; Table S5).

**Fig. 2.**
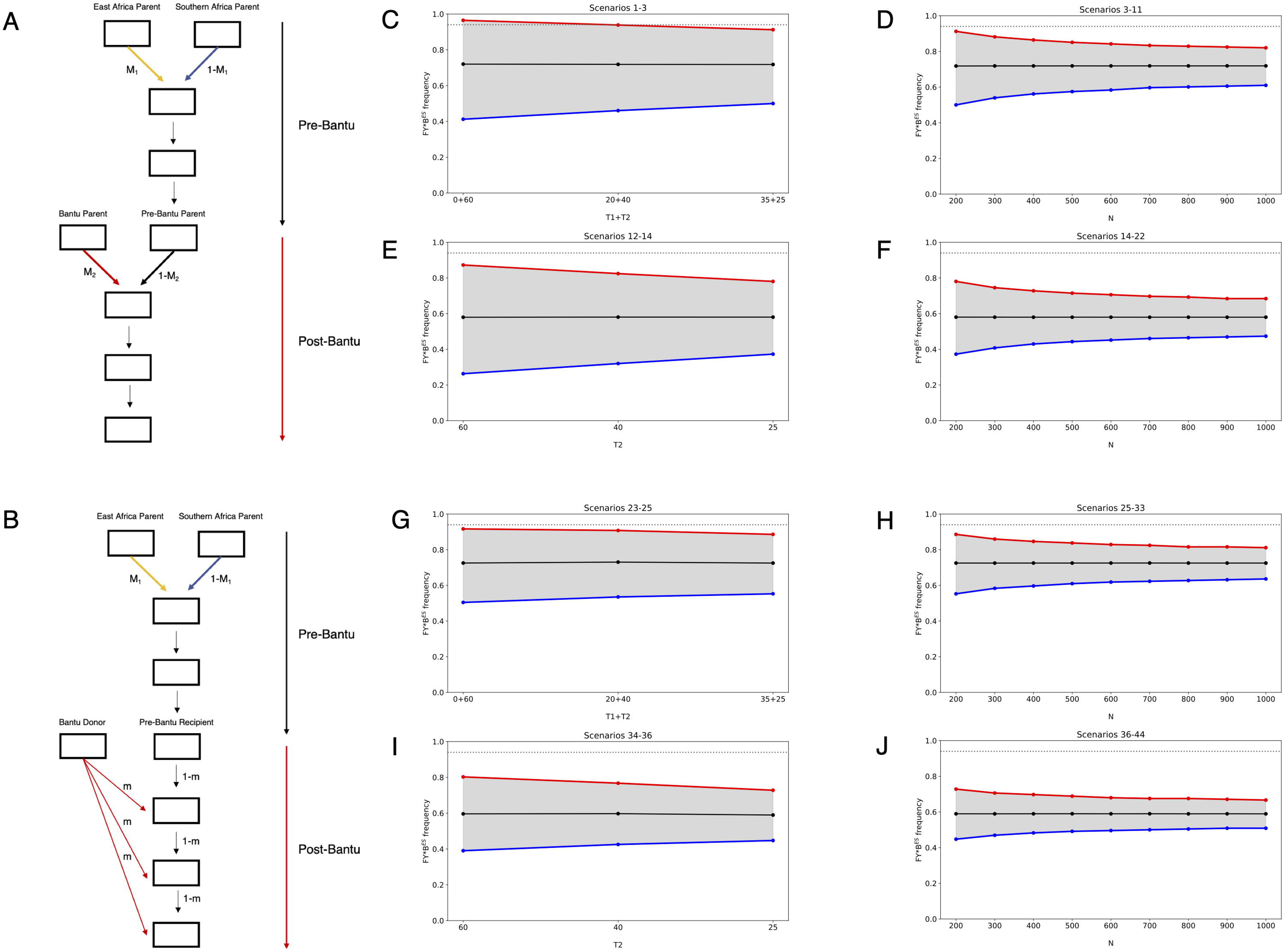
Simulations of admixture and genetic drift. Admixture models with two admixture pulses (A) and with one pulse followed by continuous gene flow (B). M1=East-African-related ancestry proportion; M2=Bantu-related ancestry proportion; m=migration rate from the Bantu donor population per generation; T1=number of generations between the first admixture pulse and the onset of Bantu admixture; T2=number of generations from Bantu admixture to present. (C-J) Distributions of simulated FY*B^ES^ allele frequencies in samples of 114 individuals, under different admixture models, with varying time intervals since admixture (T1 and T2), initial allele frequencies and population sizes (N), corresponding to scenarios outlined in Table S5: panels C-F refer to scenarios 1 to 22, and panels G-J refer to scenarios 23-44. In each panel, the dashed grey line indicates the observed allele frequency in the Khwe; the solid black line indicates the expected allele frequency; and the grey shaded area represents the 95% quantile of simulated allele frequencies.

Bantu ancestry was modelled as a second discrete admixture pulse (Fig. 2A) or as continuous gene flow into the pre-Bantu population (Fig. 2B), consistent with the spread of Khwe individuals along the PC1 axis in Fig. 1B. Each admixture scenario was tested using two different sets of FY*B^ES^ allele frequencies in the parental populations: a conservative set based on current frequencies in Ju□’hoan (11%) and Somali (92%) (Fig. 2 C-D and G-H), and a less conservative set assuming that the FY*B^ES^ allele was introduced solely via Bantu-speaking groups (Fig. 2 E-F and I-J). All simulations used a previously estimated date of 60 generations for admixture between southern and eastern African populations [25], while the timing of Bantu admixture was allowed to vary between 60 and 25 generations ago, spanning the upper range of estimates for admixture between Bantu speakers and pre-Bantu groups in southern Africa [25,26,43,44,54].

Each admixture scenario and set of parental allele frequencies was further combined with varying effective population sizes (Fig. 2 D, F, H, J) to assess the probability of reaching the observed FY*B^ES^ frequency of 94% by drift alone. We conducted 100,000 replicates for each demographic configuration.

Except for a single highly conservative condition involving early Bantu admixture, small effective population sizes, and high initial FY*B^ES^ frequencies in the pre-Bantu hybrid population (Fig. 2C), all tested scenarios resulted in fewer than 5% of simulations reaching or exceeding the observed 94% frequency (Fig. 2; Table S5).

These findings indicate that genetic drift alone is unlikely to account for the high FY*B^ES^ frequency in the Khwe, supporting a role for positive selection acting on the allele after admixture.

### 3.4 Ancestry deviations at the Duffy locus

After admixture, the ancestry of a source population carrying a beneficial allele is expected to become overrepresented in the genomic regions surrounding the advantageous variant [40,55].

To assess whether genomic segments containing the Duffy locus showed an excess of Bantu-related ancestry in the Khwe genomes, we inferred local ancestry while excluding the SNPs that determine Duffy allelic variation from the analysis.

We identified a 390 kb region on chromosome 1, encompassing the Duffy gene, where Bantu-related ancestry was significantly elevated (75%) compared to the genome-wide average (58%). This difference corresponds to 2.68 SD above the mean, placing the region in the 99th percentile of the genome-wide distribution of Bantu ancestry (Fig. 3A and D; Table S6).

**Fig. 3.**
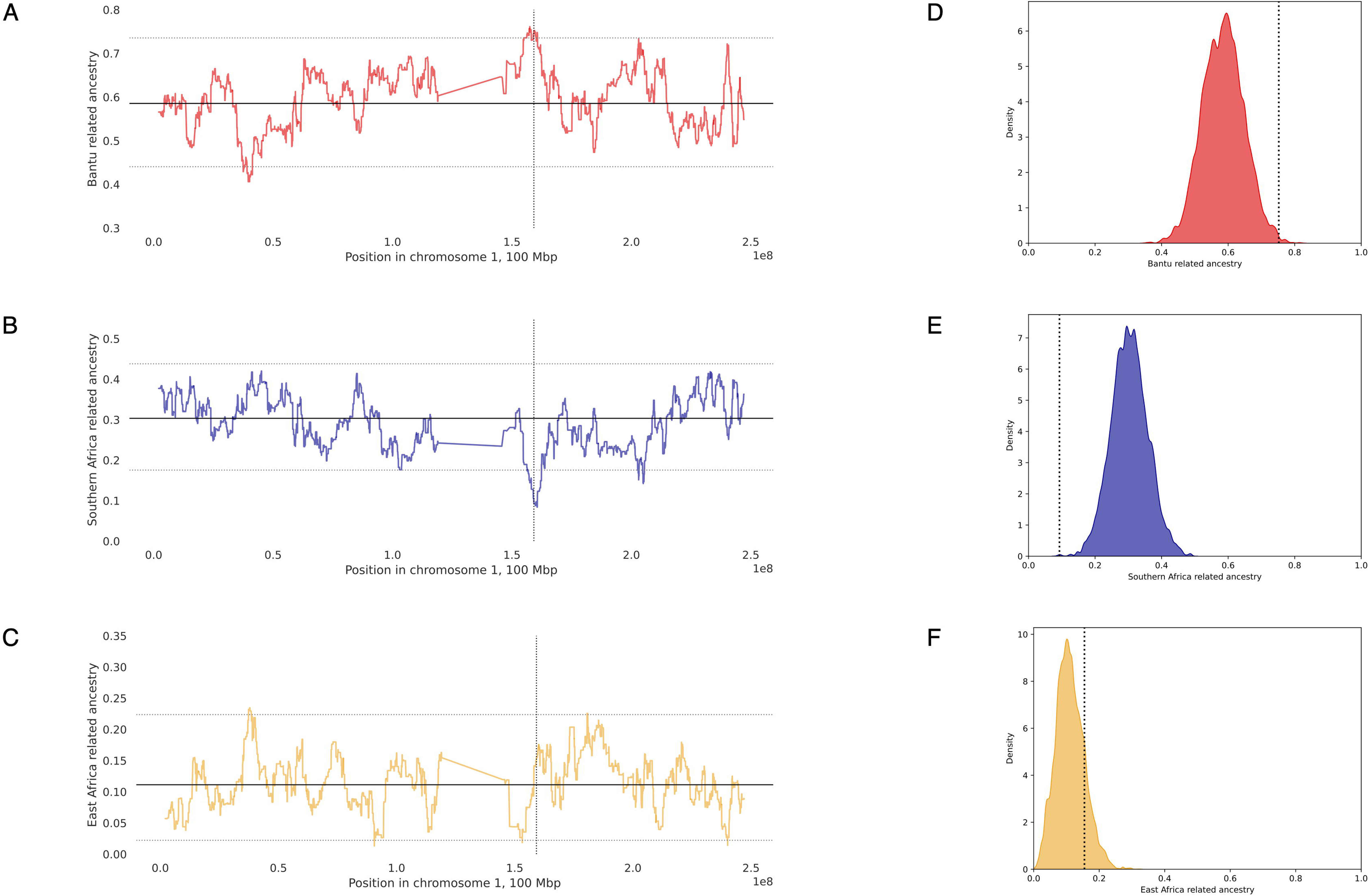
(A-C) Average local ancestry proportions in 114 Khwe individuals along chromosome 1 for Bantu (A), southern African (B), and eastern African-related (C) ancestries. Solid horizontal lines indicate genome-wide average ancestry proportions, while dashed horizontal lines correspond to the 99th and 1st percentiles of local ancestry distributions. Dashed vertical lines mark the position of the Duffy locus. (D–F) Genome-wide distributions of average Bantu (D), southern African (E), and eastern African-related (F) ancestry proportions. Dashed vertical lines indicate the local ancestry estimate for the RFMix window containing the Duffy locus. Kwangali, Ju□’hoan and Somali were used as proxies for Bantu, southern African and eastern African-related ancestries, respectively.

An opposite and stronger deviation was observed in the same genomic region for resident southern African-related ancestry (9,3 vs 30, 5%), corresponding to 3.79 SD below the mean (Fig. 3B and E; Table S6).

The asymmetry between the two ancestry deviations appears to be driven by a non-significant local elevation of East African ancestry (Fig. 3C and F; Table S6), which could reflect random noise or the presence of FY*B^ES^ linked markers in the Somali population. When the Hadza hunter-gatherers, who share partial ancestry with the Somali [48], are used as proxies for the eastern African contribution, the local signals for elevated Bantu ancestry and decreased southern African ancestry become more symmetrical (Fig. S5). Both signals remain robust across different admixture times and phasing methods (Table S6).

Taken together, these results support the hypothesis that the FY*B^ES^ allele and its linked haplotypes underwent post-admixture positive selection in the Khwe, following the arrival of Bantu-speaking populations in the lower Okavango River Basin.

### 3.5 Strength of selection

Finally, we applied a deterministic model assuming an infinite population size to estimate the selection coefficient (*s*) required to reach the observed FY*B^ES^ allele frequency. This analysis was based on the admixture scenario shown in Fig. 2A, using different sets of initial allele frequencies and admixture times (Fig. S6; Table S7).

Assuming that selection acts only on individuals homozygous for FY*B^ES^ (*h* = 1), the estimated selection coefficients ranged from 0.035 to 0.114. Under an additive model (*h* = 0.5), the estimated *s* values ranged from 0.059 to 0.179 (Fig. S6; Table S7). These estimates are comparable to those reported in previous admixture studies in Madagascar and Cape Verde [21,22,56], suggesting that strong recent selection favors Duffy negativity in different epidemiological and ecological contexts.

## 4. Discussion

The evolutionary relationship between Duffy-negativity and *Plasmodium vivax* has been extensively debated. Several lines of evidence have been invoked to support the hypothesis that the parasite was not the sole driver of Duffy-negativity. These include the relatively mild clinical course of *vivax* malaria compared to *P. falciparum* infection, the documented ability of *P. vivax* to infect Duffy-negative individuals, and the absence of widespread *P. vivax*–resistance variants in human populations historically exposed to the parasite across Eurasia [13,57–59].

On the other hand, recent studies suggest that the burden of *P. vivax* malaria has been historically underestimated, and available data indicate that Duffy-negativity continues to confer significant protection against infection [14,15,60]. Moreover, the discovery in Papua New Guinea of the rs2814778 T>C promoter mutation on the background of the FY*A allele suggests that parallel adaptation may be occurring in regions with high *P. vivax* endemicity [61]. Finally, a role of *P. vivax* resistance in driving the high frequency of the FY*B^ES^ allele is supported by evidence that post-admixture selection favours Duffy-negativity in regions where *P. vivax* prevalence is known to be historically high, but not in areas like southern Africa, where the parasite was long thought to be absent or only present at residual levels [40].

To our knowledge, the present study provides the first evidence of post-admixture selection favouring Duffy-negativity in the lower Okavango Basin—a region of southern Africa where *P. vivax* was previously undetected but has recently been identified through highly sensitive molecular techniques, primarily in Duffy-negative individuals [23,24,62,63].

The implications of this finding critically depend on how long *P. vivax* has been present in the region. If the parasite lineages currently circulating among Duffy-negative individuals in Africa were newly adapted strains that were only recently introduced in the Okavango Basin, our results could suggest that *P. vivax* is not a selective force driving the rise in Duffy-negativity. However, since available evidence shows no clear genetic differences between strains infecting Duffy-negative and Duffy-positive individuals [64,65], it remains plausible that the circulating lineages are not newly emerged, but rather remnants of older parasite populations that were only partially suppressed by the rise in FY*B^ES^ frequency [66].

In this context, our observation that post-admixture selection has promoted the diffusion of the beneficial FY*B^ES^ allele from ancestral Bantu-speaking groups into the Khwe suggests that *P. vivax* could be circulating in the region at least 500 to 1,000 years ago, as implied by available estimates of the timing of admixture between Bantu incomers and the Khwe, as well as other pre-Bantu populations in southern Africa [25,26,43,44,54].

Due to regional variation in temperature and aridity, the historical spatial distribution of *P. vivax* endemicity in southern Africa was likely highly heterogeneous. The areas of the Okavango Basin inhabited by the Khwe fall within the southernmost ecologically defined margin suitable for stable *P. vivax* transmission in Africa [67,68]. However, available evidence suggests a broader historical distribution of *P. vivax* and associated signals of post-admixture selection.

While in present-day Botswana, the Okavango Delta and surrounding regions remain important foci of malaria transmission, recent active surveillance has also reported unexpectedly high *P. vivax* prevalence in southern districts [24]. Moreover, the frequency of the FY*B^ES^ allele (87.5%) observed in earlier serological studies among Khoe-Kwadi– speaking Shua individuals from the Nata River Valley [69] is substantially higher than expected based on their genome-wide Bantu ancestry estimates (43–49%) [25,26,48], suggesting that post admixture selection for Duffy negativity may also have occurred in this region.

In contrast, no evidence of post-admixture selection has been detected in other southern African populations, despite their contact with Bantu-speaking groups carrying the FY*B^ES^ allele [37,52,70]. This geographic contrast may reflect spatial differences in malaria exposure and environmental suitability for *P. vivax* transmission.

Further studies on post-admixture selection in resident groups that have interacted with populations from different streams of the Bantu expansions, across diverse ecological zones, will help refine our understanding of the spatial and temporal dynamics of the relationship between Duffy-negativity and *P. vivax* malaria in southern Africa.

## Acknowledgements

This study would not have been possible without the administrative and logistic support of the TwinLabs established between BIOPOLIS-CIBIO and the Universities of Namibia and Botswana. We further extend our gratitude to the Ministries of Health of the governments of Namibia and Botswana for granting us research permits, as well as to the local authorities and institutions that supported our fieldwork. We specifically wish to acknowledge the Teemashani Trust representing the □Ani-Khwe community at Mohembo (North-West District, Botswana), as well as Thaddeus Chedau, Mmoloki Mogomotsi, and Kelatlhilwe Gamaxo Moses who assisted us during the data collection. Finally we thank Mark Stoneking for comments and suggestions on a previous version of this manuscript.

## Author contributions: CRediT

**Artem Kasianov:** Formal analysis; Methodology; Software; Visualization

**Anne-Maria Fehn:** Conceptualization; Investigation; Visualization

**Maitseo Bolaane:** Conceptualization; Investigation

**Gaseitsiwe Masunga:** Conceptualization; Investigation

**Ezekiel Chimbioputo Fabiano:** Conceptualization; Investigation

**Jorge Rocha:** Conceptualization; Investigation; Funding acquisition; Methodology; Project administration; Writing – original draft

## Data availability

The genotype data from 114 Khwe individuals and 35 Kwangali individuals will be made available in the European Genome-Phenome Archive (EGA), and an accession number will be provided. The code is available on GitHub (https://github.com/ArtemKasianov/DuffyPaper)

## Declaration of generative AI and AI-assisted technologies in the manuscript preparation process

During the preparation of this work the authors used ChatGPT in order to improve language and readability. After using this tool, the authors reviewed and edited the content as needed and take full responsibility for the content of the published article.

