## Supplementary materials for "Post-admixture selection favours Duffy negativity in the Lower Okavango Basin"

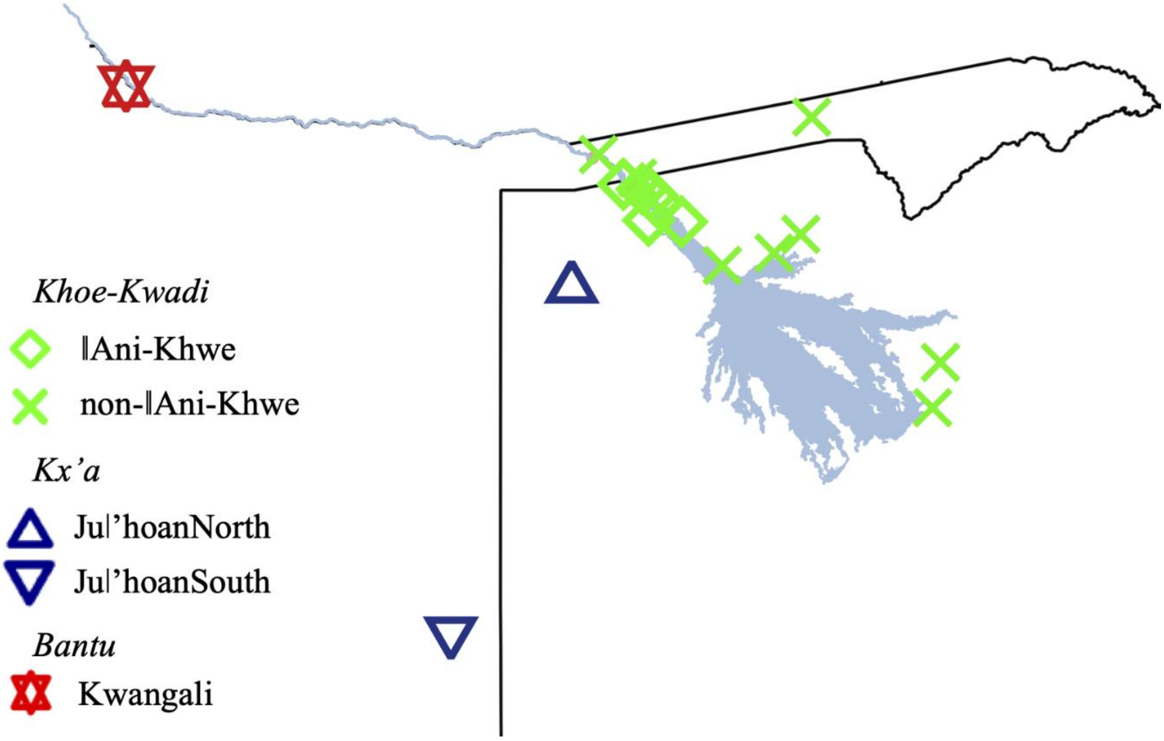


**Fig. S1**. Geographic locations of sampled Khwe-speaking individuals, together with Kwangali (this study) and Ju|'hoan-speaking groups [1,2], which were used as proxies for parental populations.

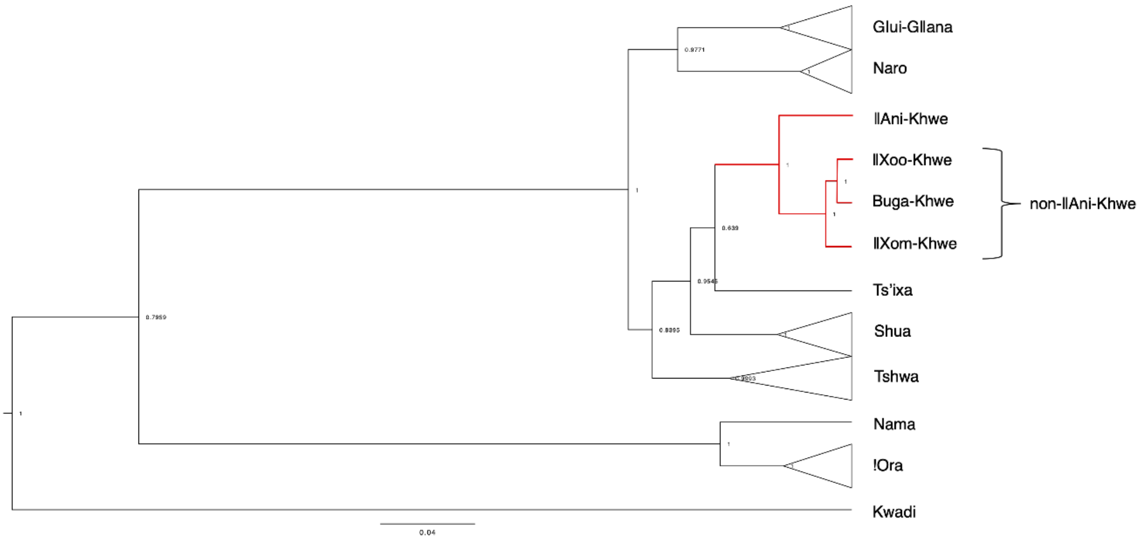


**Fig. S2**. Consensus tree of a Bayesian phylogenetic analysis of the Khoe-Kwadi language family under the Continuous Markov Chain Model. The numbers indicate posterior probabilities for each split. Branches corresponding to different Khwe dialects are marked in red (modified from [3]).

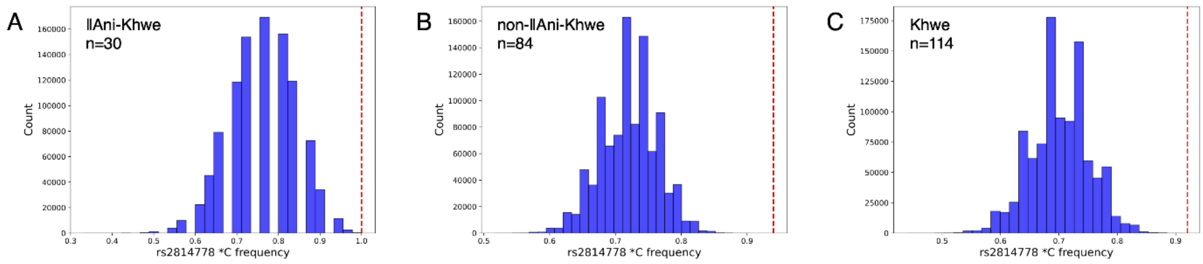


**Fig. S3.** Distributions of expected FY*B$^{ES}$ (rs2814558*C) allele frequencies obtained by binomial resampling with $10^6$ replicates. The dotted lines indicate the observed allele frequencies in each Khwe subgroup (A, B) and in the total Khwe sample (C).

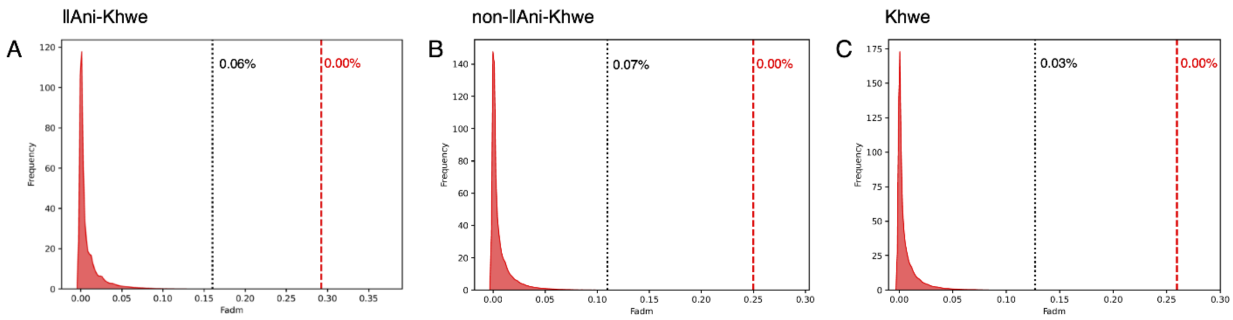


**Fig. S4.** Genome-wide distributions of the $F_{adm}$ statistic. The vertical lines indicate the $F_{adm}$ values for the SNP rs2814778 in each Khwe subgroup (A, B) and in the total Khwe sample (C). Dotted black lines indicate $F_{adm}$ values calculated with the most conservative array of parental rs2814778*C (**(**FY*B$^{ES}$) allele frequences: Kwangali 100%; Juǀ'hoan 11%; Somali 92%. Dashed red lines indicate $F_{adm}$ values assuming that the FY*B$^{ES}$ allele was absent in the pre-Bantu hybrid population. The numbers refer to the percentages of all SNPs exhibiting values equal or greater than that of the rs2814778 SNP under each assumption.

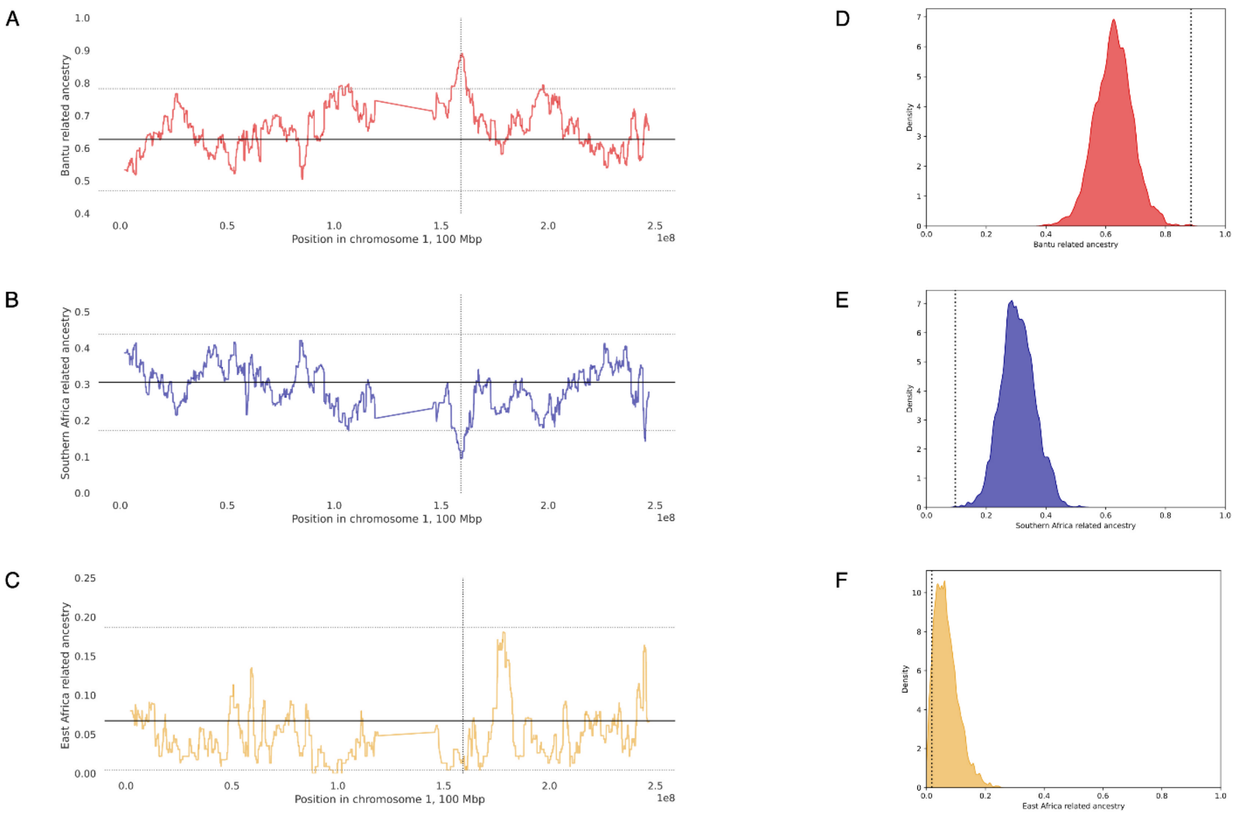


**Fig. S5.** (A-C) Average local ancestry proportions in 114 Khwe individuals along chromosome 1 for Bantu (A), southern African (B), and eastern African-related (C) ancestries. Solid horizontal lines indicate genome-wide average ancestry proportions, while dashed horizontal lines correspond to the 99th and 1st percentiles of local ancestry distributions. Dashed vertical lines mark the position of the Duffy locus. (D–F) Genome-wide distributions of average Bantu (D), southern African (E), and eastern African-related (F) ancestry proportions. Dashed vertical lines indicate the local ancestry estimate for the RFMix window containing the Duffy locus. Kwangali, Ju|'hoan and Hadza were used as proxies for Bantu, southern African and eastern African-related ancestries, respectively.

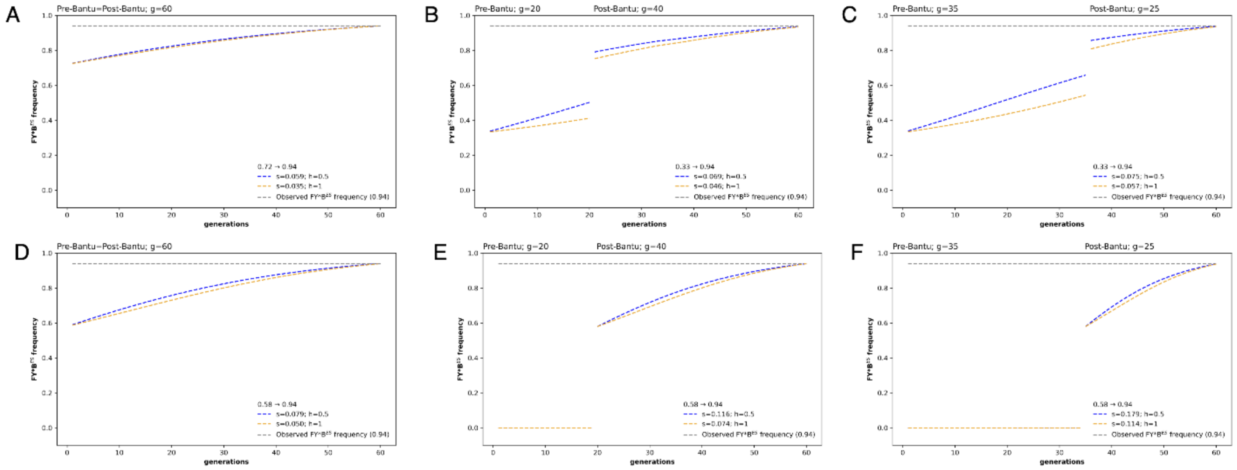


**Fig. S6.** Trajectories of FY*B$^{ES}$ allele frequencies under the admixture scenarios described in Table S7, using the best-fitting selection coefficients ($s$), dominance coefficients ($h$) of 0.5 and 1, and assuming an infinite population size.
